# Structural decay of poly(ethylene terephthalate) by enzymatic degradation

**DOI:** 10.1101/2025.03.24.645130

**Authors:** Daisuke Tadokoro, Tomoya Imai

**Affiliations:** Research Institute for Sustainable Humanosphere, Kyoto University, Gokasho, Uji, Kyoto 611-0011, Japan

**Keywords:** poly(ethylene terephthalate) (PET), PETase, plastic recycling, enzymatic degradation, porous

## Abstract

Synthetic polymers, such as plastics, are ubiquitous materials that are used in many applications. The sustainable use of plastics is becoming increasingly important given the emergent issues of environmental pollution by microplastics and the limited resources of petroleum on Earth. One of the key strategies for the sustainable use of plastics is recycling. Enzymatic degradation is a promising technique for recycling plastic because this process requires neither energy nor harsh solvents, such as strong alkaline solutions and organic solvents. In this study, the enzymatic degradation of poly(ethylene terephthalate) (PET), a major plastic used in daily life, was investigated, aiming to improve the efficiency of enzymatic degradation by understanding the decay of the polymeric PET structure. The structural decay of an amorphous PET film induced by a PET-hydrolyzing enzyme (PETase) was analyzed using wide-angle X-ray diffraction (WAXD), small-angle X-ray scattering (SAXS), electron microscopy, and X-ray computed tomography (X-ray CT). Structural decay progressed from the surface of the film, and many nested pores (10^-8^‒10^-5^ m) were found in the later stage of degradation, reflecting the structural difference between the surface and core of the material. Information regarding the polymeric structure of the plastic under enzymatic degradation is important for improving the degradation performance of enzymes for realizing the biochemical recycling of plastics such as PET.

**Highlights:**

- PETase-induced decay of a PET film was analyzed regarding its polymer structure
- Decay occurred from the surface to form multiscale pores of 10^-5^ to 10^-8^ m
- The PET decay mechanism differs from that of cellulose, a typical biopolymer
- Cellulose biodegradation is a good example to learn biodegradable PET materials

## 1. Introduction

Synthetic polymers and plastics are important materials in our daily lives. Massive amounts of plastics are produced from petroleum, which is a limited resource. Accordingly, securing a supply of raw materials that can replace petroleum or naphtha is becoming increasingly important for sustainable life. Another issue with the use of plastics is environmental pollution. Ocean plastics are a major environmental pollutant and social issue owing to their significant threat to marine organisms [1, 2]. The impact on human health has also been debated [3]. Therefore, conserving natural resources and reducing the emission of environmental pollutants are important for the sustainable use of plastics in the future.

Thus, a circular plastic economy must be realized [4, 5] to secure raw materials for regenerating plastics and reduce the leakage of plastics into the environment. Poly(ethylene terephthalate) (PET) is a popular plastic that is widely used in food packaging, including drink bottles and cost-effective clothing. In Japan, PET is the third most common marine plastic waste in coastal waters, after polyethylene and polypropylene [6], and the situation is expected to be similar in other countries.

There are two types of PET recycling systems: mechanical and chemical [7]. In the mechanical recycling of PET, PET materials such as drink bottles are reshaped by heating after fragmentation into flakes. This process is relatively simple and is the primary method for recycling PET. However, the process of mechanically crushing the material into smaller flakes for efficient thermal reshaping is a potential origin of microplastics in the environment and a possible issue in achieving a highly sustainable society with plastics. In contrast, for chemical recycling, the used PET material is first disintegrated into monomers or oligomers via chemical reactions. The obtained monomers and oligomers are recovered and polymerized to regenerate PET. This process can maximize the recycling efficiency because it can be applied to various materials (colored bottles, yarns, textiles, films, etc.). However, the process is costly because of the energy and chemicals required to disintegrate the used PET materials and recover the monomers and oligomers.

Recently, enzymatic degradation has attracted considerable attention as an efficient tool for disintegrating PET materials under mild conditions for chemical recycling. After the initial discovery of a PET-hydrolyzing enzyme in a biomass-degrading bacterium *Thermobifida fusca* [8], many studies using life-science strategies have reported PET-hydrolyzing enzymes, hereafter called PETase, based on bacterial screening [9], metagenomic screening [10], and rational design by site-directed mutagenesis [11, 12]. The mechanism of enzymatic hydrolysis of PET molecules is well debated and deeply understood [13–15]. These studies have provided impetus for improving the enzymatic activity of PETase using life-science strategies; however, only basic characterizations, such as analysis of the weight loss, surface morphology, and crystallinity using differential scanning calorimetry (DSC), have been performed [9, 11], and little attention has been paid to the decay of the higher structures of PET by PETase.

Given the current situation, the authors propose that the structure of PET materials is an important factor that governs the enzymatic degradability and properties of PET; thus, in-depth analysis of the structural changes of PET materials during degradation by PETase is warranted. In this study, a multiscale analysis of the structural decay of an amorphous PET film by PETase was conducted using Fast-PETase [11] as a model enzyme, which is an improved variant of IsPETase from a bacterium *Ideonera sakaiensis* (discovered at a PET bottle recycling site) [9], by machine-learning. The model PET substrate used in this study is a lab-made, amorphous film that has been intensively analyzed in polymer science [16–18]. Using this experimental setup, this study aims to clarify how PETase disintegrates the polymer structure of PET.

## 2. Materials and Methods

All chemicals used in this study were purchased from Fujifilm Wako Chemicals (Tokyo, Japan), Nacalai Tesque Inc. (Kyoto, Japan), and Sigma-Aldrich Inc. (St. Louis, MO, US), unless otherwise indicated. Tryptone and yeast extracts for the bacterial cultures were purchased from Becton Dickinson (Franklin Lakes, NJ, USA). Imidazole for protein purification was purchased from Calbiochem, Inc. (Rahway, NJ, USA).

### 2.1. Preparation of PET substrate

PET pellets were purchased from Scientific Polymer Products Inc. (NY, USA; catalog #: 138; inherent viscosity: 0.58) and subjected to melt-quenching to prepare amorphous PET films. Firstly, approximately 60 mg of the PET pellets were sandwiched between a pair comprising Kapton film, glass slides, and copper plates; the Kapton film touched the PET pellets directly, and the copper plates exposed to the outside were used for efficient heat transfer. This sandwich was placed on a heating plate and maintained at 300 °C for *ca.* 2 min and manually pressed with forceps to remove visible air bubbles. Thereafter, the sandwich of PET in the Kapton film was quickly placed between the copper plates at room temperature (*ca.* 20 °C), which is lower than the glass transition temperature of PET (*ca.* 75 °C). This melt-quenched amorphous film, which typically had a thickness of 150 µm and a round shape with a radius of 10 mm, was evenly cut with scissors into 8 fan-shaped pieces and used as the substrate in the degradation experiment.

### 2.2. Purification of PETase

#### 2.2.1. Cloning of Fast-PETase

The PETase used in this study is a previously reported Fast-PETase [11], which is a highly thermostable variant of IsPETase. The first 27 amino acid residues were truncated as previously described [11] because this region was predicted to be a signal peptide by SignalP6.0 [19]. The DNA sequence of Fast-PETase without the signal peptide sequence was synthesized by Eurofins Genomics Inc. (Tokyo, Japan) with codon optimization in *Escherichia coli*. The DNA of the codon-optimized Fast-PETase gene was inserted between the NdeI and BamHI sites in the cloning site of the pET-28b plasmid (Millipore Inc.) by restriction enzyme treatment and subsequent ligation.

#### 2.2.2. Expression of recombinant PETase protein in *Escherichia coli*

The pET-28b plasmid carrying Fast-PETase, prepared as described in Section 2.2.1, was introduced into *Escherichia coli* BL21(DE3). The *E. coli* transformants carrying this plasmid DNA were selected based on kanamycin resistance on LB agar plate, and used for expressing recombinant Fast-PETase protein.

The transformant was precultured in LB liquid medium with 50 μg/mL kanamycin at 37 °C with orbital shaking at 200 rpm. In a typical experiment, 0.5 mL of the preculture was transferred into 50 mL of fresh 2× YT medium with 25 μg/mL kanamycin in a 300 mL flask without baffles. The flask was maintained at 37 °C in an orbital shaker with agitation at 200 rpm until the *OD*_600_ reached 0.6‒0.8, after which isopropyl thio-β-D-galactoside (IPTG) was added to a final concentration of 0.5 mM to induce protein expression. The culture was then maintained at 20 °C for *ca.* 20 h in an orbital shaker at 200 rpm. Finally, the cultured *E. coli* cells were collected by centrifugation at 5,000×*g* for 15 min at 4 °C and kept in a freezer at −80 °C until use.

The frozen cells were thawed on ice and suspended in phosphate buffered saline (PBS) containing 1 mM phenylmethylsulfonyl fluoride and 5–10 mM imidazole. The cell suspension was ultrasonicated to disrupt the cells, centrifuged at 12,000×*g* for 30 min at 4 °C to remove the unbroken cells, and the supernatant was filtered using Miracloth (Merck Inc.). Hexahistidine-tagged Fast-PETase in the clarified lysate was purified using immobilized metal affinity chromatography (IMAC). Ni-Sepharose (Cytiva, Inc. Marlborough, MA, US) was placed in a plastic column (Poly-Prep column; Bio-Rad Inc., Hercules, CA, US) and equilibrated with 5–10 mM imidazole in PBS. The clarified lysate was then applied to Ni-Sepharose in the column to bind the tagged Fast-PETase. The Ni-Sepharose was then washed with PBS containing 20 mM imidazole, and the tagged Fast-PETase was eluted with PBS containing 300 mM imidazole. IMAC-purified Fast-PETase was applied to BioGel P-4 or P-6 resin (BioRad Inc.) in a spin column and centrifuged at 1,000×*g* to exchange the buffer with PBS (without imidazole). The concentration of Fast-PETase was estimated from the absorbance at 280 nm, using an extinction coefficient of 1.4 (mg/ml)^-1^·cm^-1^.

### 2.3. Enzymatic treatment of PET

The fast-PETase purified as described in Section 2.2 was reconstituted at a concentration of 500 nM in 100 mM HEPES-NaOH buffer (pH8.0), and the amorphous film prepared as per Section 2.1 was immersed in this reaction buffer in a 1.5 mL polypropylene microtube. The reaction was conducted at 50 °C by placing the reaction microtubes in an air-phase incubator without exchanging the reaction buffer with fresh enzyme, and the PET film was removed from the reaction buffer at the desired time. The film was washed briefly with water and dried in air for at least 15 min, weighed using an electronic balance, and photographed using a digital camera. The dried PET film was subjected to structural analyses.

### 2.4. Structural analysis of degraded PET

#### 2.4.1. Scanning electron microscopy (SEM)

The sample treated with Fast-PETase was fixed on a copper stub using electron-conductive glue (DOTITE, Fujikura Kasei Co., Ltd., Tokyo, Japan); the sample was coated with platinum using a JEC-3000FC coater (JEOL Inc., Akishima, Tokyo, Japan) that was operated at 20 mA for 90 s with the rotating sample stage at a tilt angle of 45° for 45 s and then at 0° for another 45 s.

The platinum-coated samples were observed using a JSM-7800F prime (JEOL Inc.) instrument operated at an accelerating voltage of 5.0 keV and a working distance of 6 mm. Some images were captured from the side direction by fixing the sample film perpendicularly on a copper stub with a working distance of 10 mm.

#### 2.4.2. X-ray computed tomography (X-ray CT)

Micro-X-ray computed tomography (CT) was performed at BL8S2 of the Aichi Synchrotron Radiation Center (Aichi SRC, Seto, Aichi, Japan). Briefly, the sample was set on a rotational stage and illuminated with white X-rays (6–24 keV). The transmitted images were recorded with sample rotation from 0 to 360° at an interval of 0.1° using an ORCA-Flash4.0 sCMOS image sensor (Hamamatsu Photonics K.K., Hamamatsu, Shizuoka, Japan)) with a zoom lens: the pixel size corresponds to 0.65 μm^2^. Three-dimensional reconstruction of the obtained image dataset was performed at the beamline using TomoPy, which is a Python package for X-ray CT image analysis and image reconstruction [20]. Three-dimensional volumetric data were visualized using Fiji [21] and Molcer (WhiteRabit Co., Ltd., Tokyo, Japan) for visual inspection.

#### 2.4.3. Small angle X-ray scattering (SAXS) and wide angle X-ray diffraction (WAXD)

Small-angle X-ray scattering (SAXS) and wide-angle X-ray diffraction (WAXD) analyses were performed at the synchrotron facility, BL40B2, of SPring-8 (Hyogo, Japan). The wavelength of the X-rays used in this study was 1 Å and the scattering/diffraction patterns were recorded with a Pilatus3 S 2M instrument (Dectris Inc., Switzerland). The exposure times were 10 s and 5 s for SAXS and WAXD, respectively.

The scattering/diffraction patterns were circularly averaged using a beamline program and corrected for absorbance. Given the scattering vector *q* = (4π sin *θ*)/*λ*, where *θ* and *λ* are half of the scattering angle and the wavelength of the X-rays, respectively, the *q*-range of the SAXS and WAXD experiments covered 0.007‒0.4 Å^−1^ and 0.05‒2.5 Å^−1^, respectively. For some data, the WAXD profiles were manually rescaled and merged with the SAXS profiles using Microsoft Excel.

## 3. Results and Discussion

### 3.1. Weight-loss and visual inspection of amorphous PET film after Fast-PETase treatment

First, the weight loss of the amorphous PET film treated with purified Fast-PETase was evaluated (Figure 1), conforming the significant PET-degrading activity of the Fast-PETase used in this study. Visual inspection also demonstrated a change from the clear appearance of the PET film before degradation to white after degradation by Fast-PETase, where the whiteness became more pronounced after degradation (Figure 1), consistent with prior reports [9].

**Figure 1.**
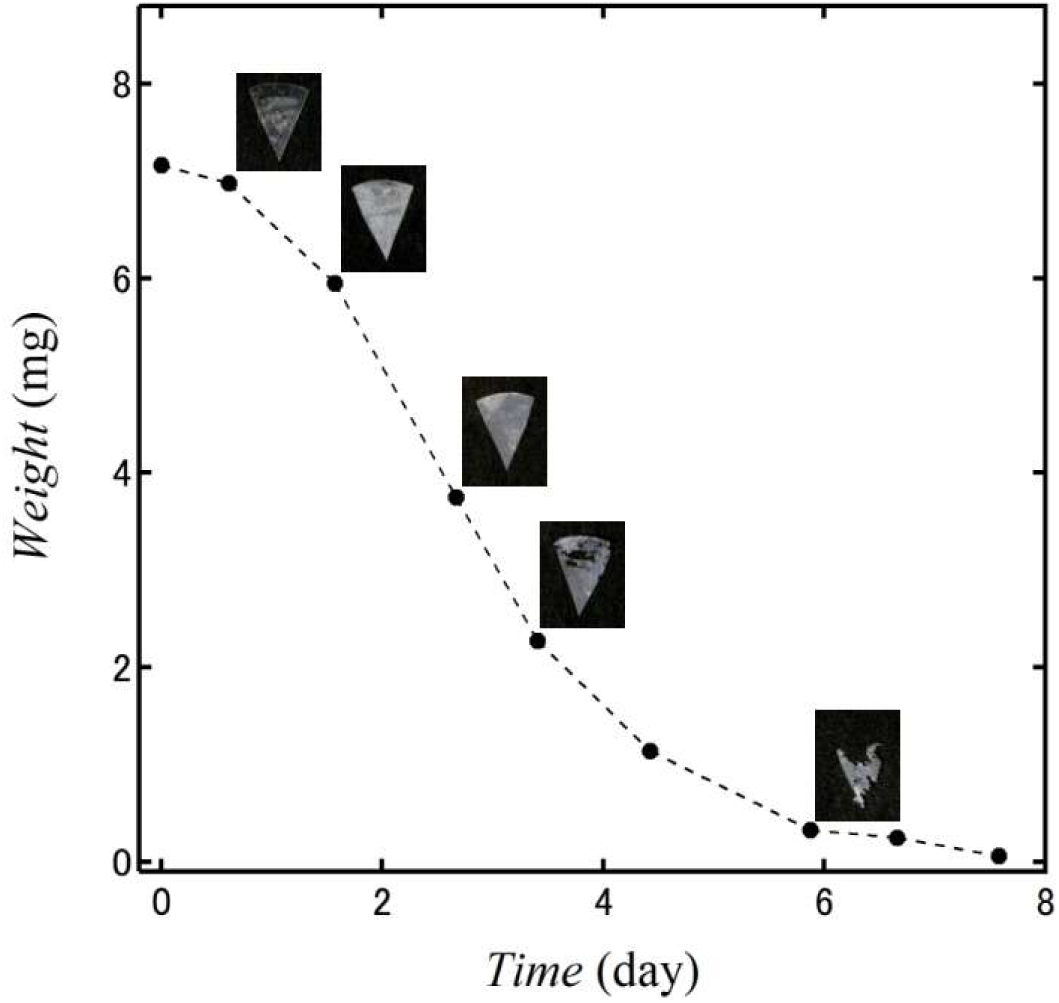
Representative weight loss data and visual appearance of PET film degraded by Fast-PETase.

Two mechanisms were hypothesized for this phenomenon: (i) an increase in the crystallinity upon degradation and (ii) the formation of a structure with voids having dimensions on the order of the wavelength of visible light (∼10^-7^ m) in the PET film sample. Given a lower glass transition temperature for the surface of a polymer film [22], crystallization might be possible under the glass transition temperature of PET (*ca.* 50 °C) and was tested as a reason for whitening in this study. The residual PET films were analyzed using WAXD, X-ray CT, SEM, and SAXS to test these hypotheses. Furthermore, the structural changes in the PET film due to enzymatic degradation were tracked over the reaction time.

### 3.2. WAXD analysis

First, the fine structure of the film was assessed using WAXD to test the hypothesis that the increased turbidity of the film induced by Fast-PETase was due to the increased crystallinity of PET. The WAXD profile of PET before degradation (Figure 2A) shows no sharp peaks, but a broad peak composed of a number of diffraction peaks, indicating that the melt-quench film is amorphous. The intensity of the broad diffraction peak decreased with the degradation time (Figure 2A), indicating that PET molecules were gradually removed from the film by Fast-PETase. However, the sharpness of the WAXD profile did not change significantly, indicating that the film remained highly amorphous after degradation. Thus, the turbidity of the PET film observed after degradation by Fast-PETase was not caused by increased crystallinity of PET.

**Figure 2.**
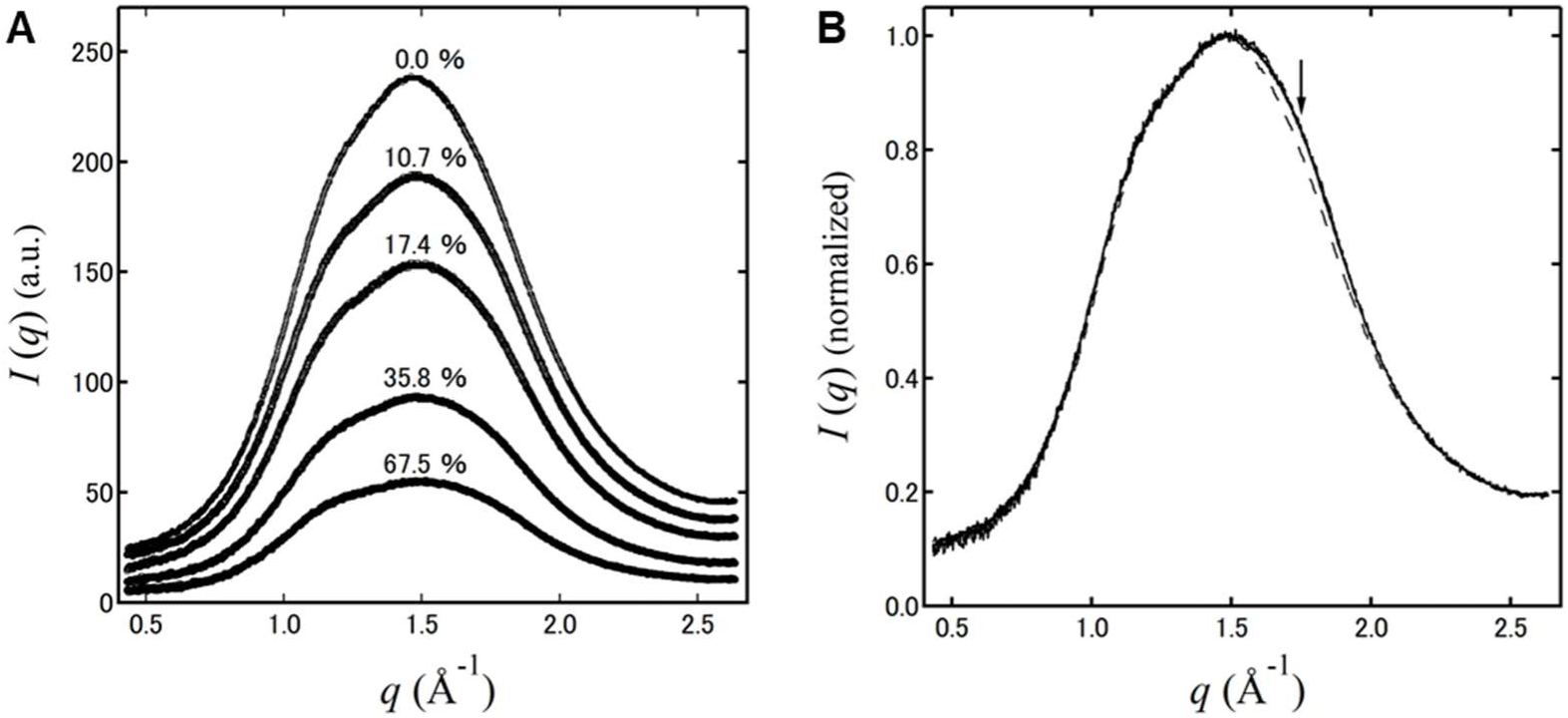
WAXD profiles of PET films degraded for different times: 0 h, 15.5 h, 26.5 h, 55.2 h, and 64.5 h. (A) Absorbance-corrected profiles with the weight loss percentage indicated for each of the profiles. (B) The WAXD profiles were normalized to the most intense peak at *ca.* 1.5 Å^-1^. Dashed line corresponds to the film before degradation, while the solid lines indicate PET degraded for 15.5 h, 26.5 h, 55.2 h, and 64.5 h; all profiles were similar. The arrow indicates the *q*-region where the diffraction intensity increased after degradation.

The small difference in the WAXD profiles at different enzymatic degradation times suggests that Fast-PETase evenly degraded the amorphous PET film on a scale observable by WAXD. In other words, Fast-PETase seems to have no preference for more crystalline PET substrates when degrading PET. This is in contrast to cellulose-degrading enzymes such as cellobiohydrolase, which degrade crystalline cellulose more effectively than endoglucanase [23]. Such variability in enzymatic properties is important for the effective degradation of native cellulose, a long crystalline fiber [24].

For detailed analysis of the change in the diffraction profile, all WAXD profiles were normalized relative to the most intense peak (Figure 2B). Notably, the profiles for degradation at 15.5, 26.5, 55.2, and 64.5 h were completely superimposable, whereas a slight increase in the diffraction intensity was observed in a limited *q*-region around 1.75 Å^-1^ between 0 h and 15.5 h (indicated by the arrow in Figure 2B). This small increase in the diffraction intensity is plausibly due to broadening of the amorphous hallow, suggesting a slight enhancement in the disorder of the PET molecules at the beginning of degradation. Given that this change was very small and occurred only at an earlier stage, the enhanced disorder due to PETase-induced degradation may have resulted from the removal of the outer surface layer of the PET film. Such a change, specific to the earlier stage of degradation, was also observed in the SEM and SAXS analyses, as shown in Sections 3.4 and 3.5, respectively.

### 3.3. Micro X-ray CT analysis

Given the WAXD results, the structural changes were analyzed on a larger scale. X-ray CT demonstrated that the surface morphology of the PET film was flat and smooth before degradation (Figure 3). During degradation, the film gradually became thinner and exhibited a rougher surface. In most parts of the film, no significant void spaces were observed (Figure 3A–D). These observations indicate that the erosion of the PET film by Fast-PETase primarily occurred at the surface.

**Figure 3.**
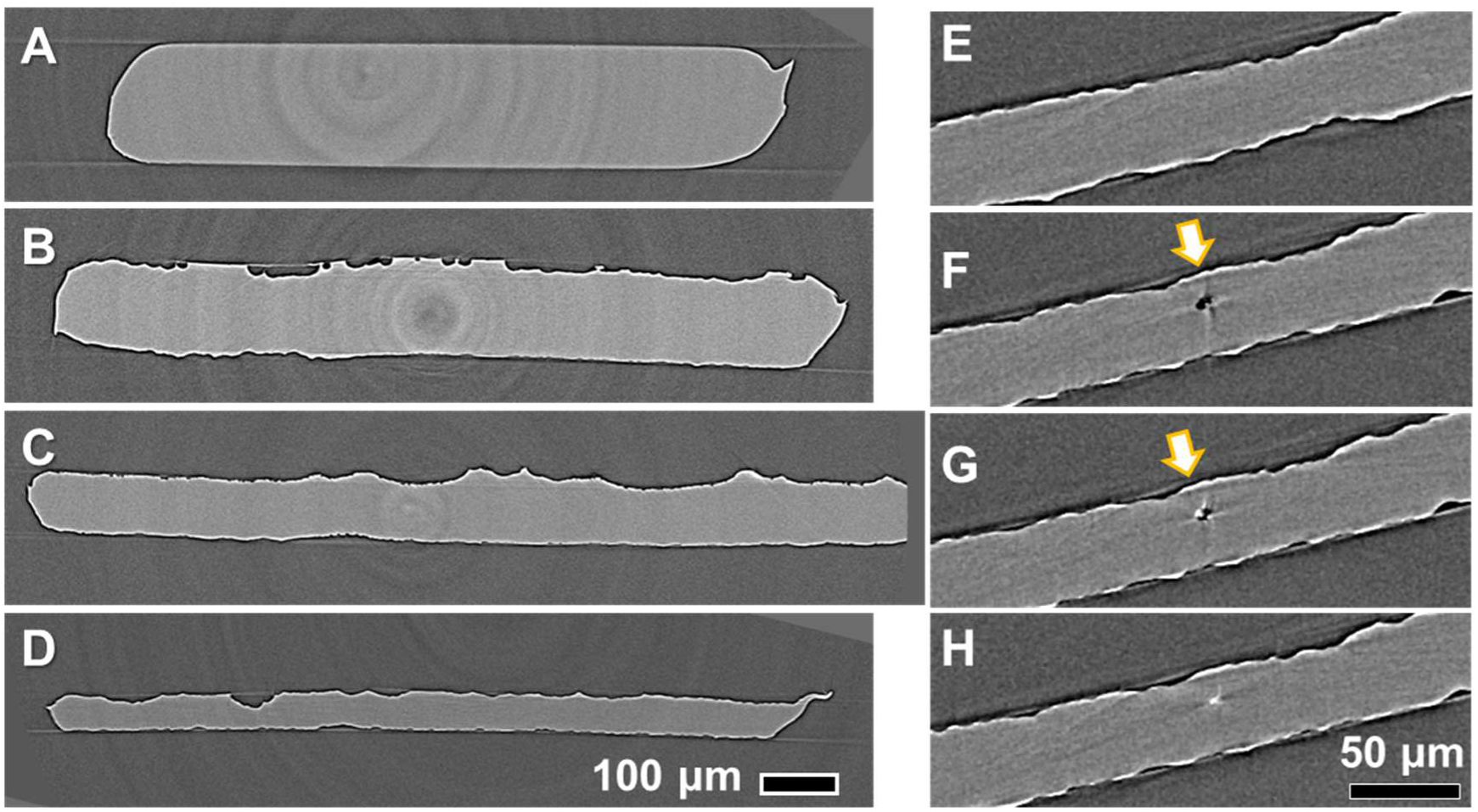
Three-dimensional volume data from X-ray CT. (A‒D) Typical cross-section of volume data for films degraded for 0 h (A), 26.5 h (B), 55.2 h (C), and 64.5 h (D). (E‒H) Serial cross-sections with an interval of 1.3 μm for the film degraded for 55.2 h.

However, using X-ray CT, some void spaces were observed in the films subjected to a longer degradation time (Figure 3E–3H), suggesting degradation of the PET film by Fast-PETase, not only on the surface but also within the film. The reconstructed X-ray CT data showed that the voids within the film were disconnected from the outside. Given that no void space was observed in the X-ray CT data before degradation of the PET film, the voids are thought to be generated by Fast-PETase. Thus, it is reasonable to assume that the pores were too small to be visualized using X-ray CT (nominally 0.65 μm pixel size): the Fast-PETase protein has a size of about 3 nm. Plausibly, the void space observed in the film using X-ray CT is not due to an experimental artifact but represents the action of Fast-PETase. Therefore, it can be concluded that gradual erosion from the surface is the major mode by which Fast-PETase attacks the PET film; however, erosion inside the film sometimes occurs at a later stage of the reaction.

Moreover, one side of the film surface became rougher than the other as the reaction progressed (Figure 4A–4D). Although this indicates a difference in the rate of structural decay of the two surfaces of the film, the reason for the difference is not clear. However, this phenomenon can be attributed to the method of preparation of the amorphous PET film. In other words, PET film preparation could be one of the keys to controlling the enzymatic digestibility, which is important for developing highly degradable PET materials.

**Figure 4.**
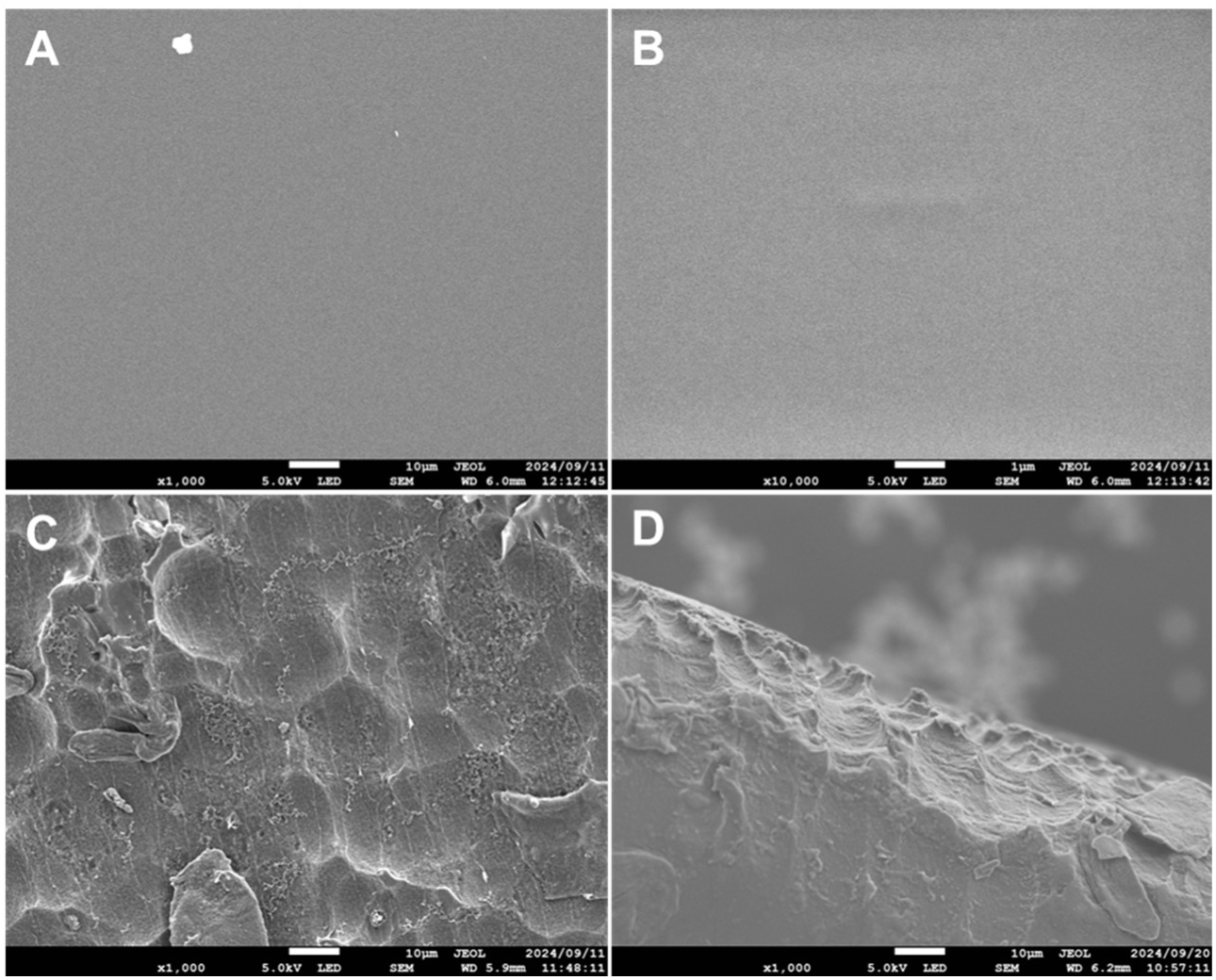
SEM images of PET film before (A and B) after degradation for 26.5 h (C and D). Scale bars indicate 10 μm except for B where it represents 1 μm. (A and B): PET before degradation at magnification of ×1,000 (A) and ×10,000 (B). (C and D): Largely degraded part of the PET film observed at a magnification of ×1,000 in top-view (C) and side-view from the edge (D).

### 3.4. SEM analysis

SEM observation in top-view clearly demonstrated that the initially very flat and smooth surface of the PET film before degradation (Figure 4A and 4B) became undulated after degradation by Fast-PETase (Figure 4C and 4D), as previously reported [9, 11]. Observations of the surface from the side confirmed the presence of depressions that were not convex (Figure 4D).

The films were further observed at different degradation times to track the structural changes in the amorphous PET film induced by Fast-PETase (Figure 5). Before degradation, the surface was flat and smooth, whereas after 4 h of degradation, many depressions having a diameter of 1 μm were apparent. Given that areas of undegraded surface were found close to these depressions, the depressions were structures generated by Fast-PETase in the initial phase. The surface of this depression was not as rough as that of the PET degraded for 12 h, as shown below.

**Figure 5.**
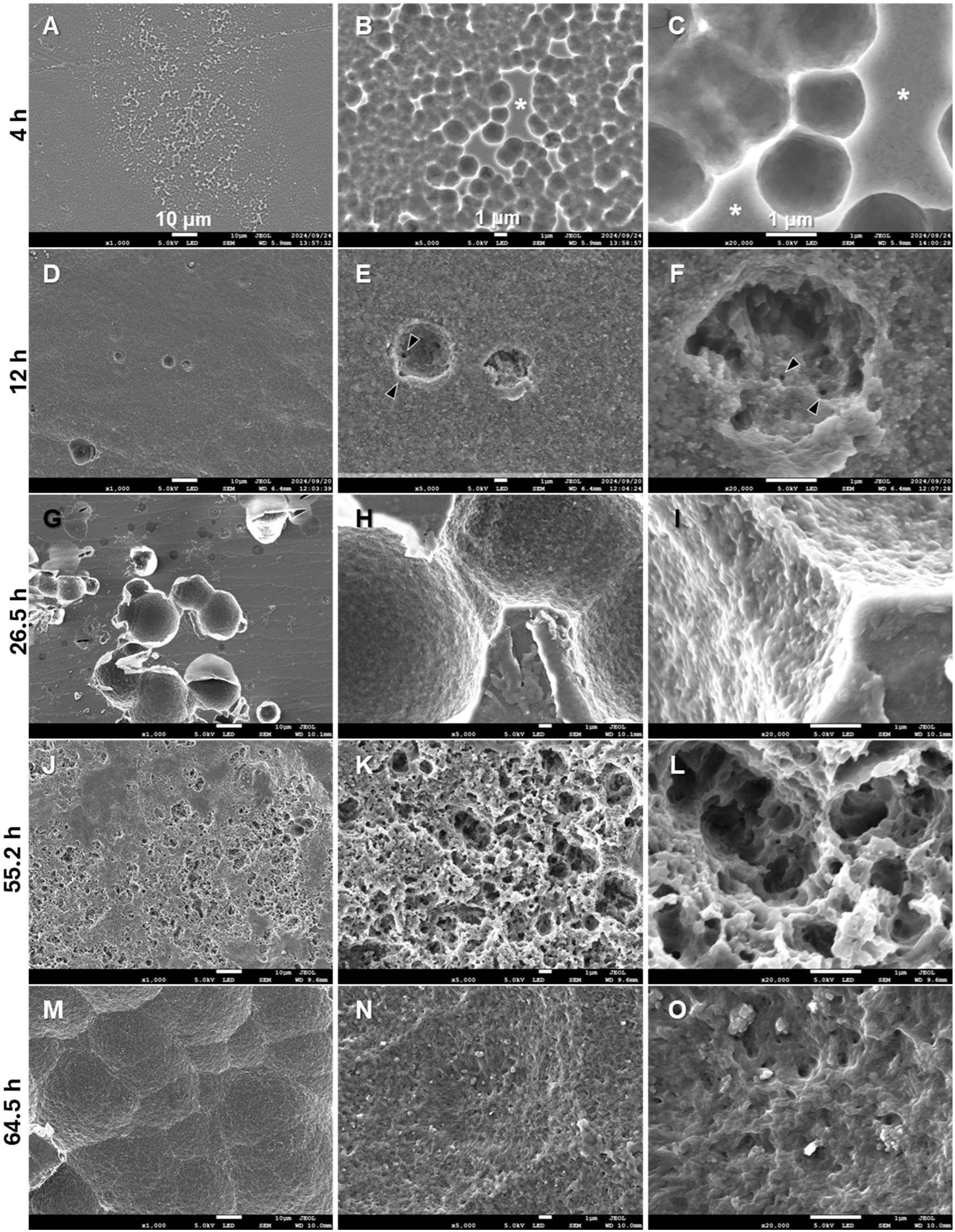
SEM observation of morphological changes of film surface during treatment with Fast-PETase at three magnifications: ×1,000, ×5,000, and ×20,000. A–C: 4 h; D–F: 12 h; G–I: 26.5 h; J–L: 55.2 h; M–O: 64.5 h. Asterisks in B and C show a flat and undegraded surface, and the arrowheads in E and F show pores with diameters of 10‒100 nm.

At 12 h of degradation, the depressions formed at 4 h were not observed, and the surface appeared flat at a lower magnification (Figure 5D). However, hollows with a diameter of 2‒4 μm appeared, and the surface appeared sandy at a higher magnification due to the granularity of many smaller pores with a diameter of 10‒100 nm (indicated by the arrowheads in Figure 5E and 5F). This hollow structure was also found less frequently at 4 h of degradation and is thus considered a characteristic feature that appears after the initial depressions (Figure 5A–5C) disappear.

After 26.5 h, the surface became highly rough even at a lower magnification due to the larger diameter (10 μm or more) of the holes (Figure 5G). Closer observation revealed many pores with diameters of 100 nm or less on the surface of the depressions (Figure 5H and 5I), which were found specifically in the hollows of the sample degraded for 12 h (Figure 5E and 5F). Thus, the hollows with sizes of a few micrometers frequently observed after 12 h of degradation might have enlarged or fused to form larger holes.

After 55.2 h, the crater-like depressions, which were also found in some areas after 26.5 h, were sufficiently enlarged to expose a relatively flat and smooth surface when viewed at lower magnification (Figure 5J). However, the higher-magnification images clearly show more porous surfaces with smaller pore sizes (10‒100 nm) and larger numbers of pores than at 26.5 h (Figure 5K and 5L). These porous features were maintained at 64.5 h, whereas the 1 μm pores became less; consequently, the surface became flatter with pores of 100 nm or less in diameter (Figure 5N and 5O). In a wider view (Figure 5M), the crater-like depressions with a diameter of ∼50 μm appeared more clearly. In summary, degradation by Fast-PETase induced hierarchical boring on the order of 10^-5^ m to 10^-8^ in the PET film.

It was thus concluded that the turbidity of the PET film after Fast-PETase degradation was due to the porous structure generated by Fast-PETase and not to crystallization of the amorphous parts, as discussed in Section 3.1.

### 3.5. SAXS analysis

Although scanning electron microscopy (SEM) is a powerful tool for visualizing the fine structure of degraded PET, it only provides a local view of the structure. Therefore, it is necessary to consider whether the SEM observations truly reflect the structure as a whole. SAXS was used to visualize the mesoscale structure of the degraded PET film in an averaged view. Two notable observations were made.

Firstly, the scattering intensity around *q* = 0.022 Å^-1^, observed before degradation, decreased quickly (in the first 12 h) after treatment with Fast-PETase (Figure 6A), indicating that the structure represented by this scattering feature was selectively attacked and removed by Fast-PETase. Furthermore, this signal shifted to lower *q*, accompanied by a decrease in the intensity during the early stages of degradation, as shown in the Lorentz-factor-corrected plot (Figure 6A, inset). Given the equation *d* = 2π/*q*, where *d* indicates the spatial periodicity, the size of the structural component represented by this scattering changed from *ca.* 290 to 320 Å. Although careful discussion is required for describing the structure represented by this scattering signal, it may represent a randomly distributed denser region in a less dense region, which has been proposed as a “nodule” in some previous studies [25, 26].

**Figure 6.**
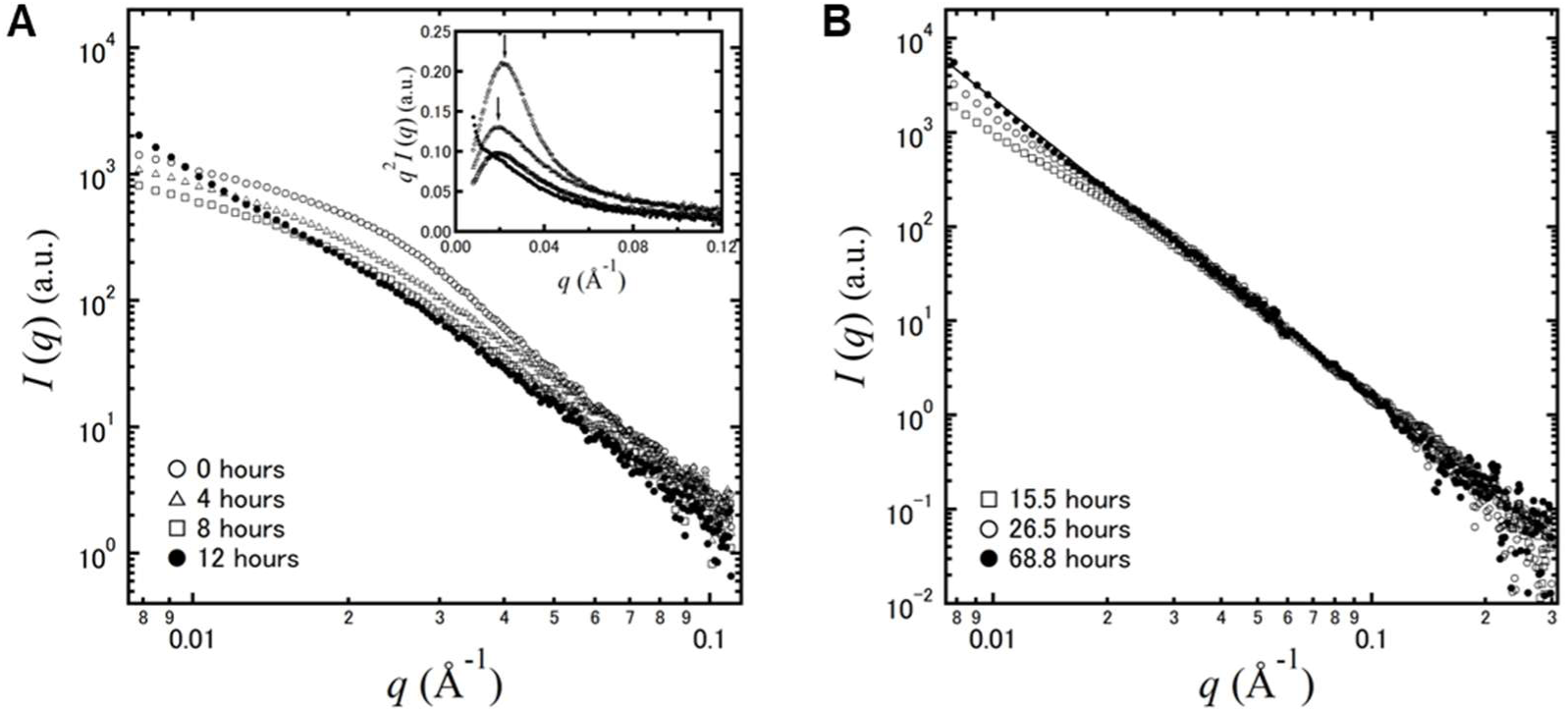
SAXS profiles of PET film treated with Fast-PETase for different reaction times. The degradation experiment was independently conducted to obtain the data in A (earlier stage of the degradation) and B (later stage). Lorentz-factor-corrected plot (*q* – *q*^2^·*I* plot) is also shown in the inset of panel A; arrows indicate the peak maxima for the films degraded for 0 h and 4 h. The scattering profiles in B were prepared by merging the WAXD data with SAXS data as described in 2.4.3. The data acquired after 68.8 h of degradation were fitted (closed circles in panel B) and followed a power-law with an exponent of −3.2 along the scattering vector *q*, as shown by black line.

The second specific structural change induced by Fast-PETase was observed as an increase in the scattering at the *q*-region below 0.02 Å^-1^ in the SAXS profile, up to the reaction time of 12 h. Given the synchronicity with the appearance of small pores in the SEM images (diameter of 10‒100 nm), this may represent the small pores observed in the SEM images after degradation for 12 h and beyond (Figure 5), where the contrast between the PET substrate and air becomes more pronounced due to the boring. The scattering in this low *q*-region increased continuously after 12 h of degradation, and finally the scattering profile at 68.8 h followed a power law with an exponent of −3.2 in the observed *q*-range (Figure 6B, black line), which provides the surface fractal dimension of 2.8 (= 6 – 3.2), indicating that the pores with diameters of 10^-7^‒10^-8^ m formed a three-dimensionally rough surface rather than a smooth surface. This is also consistent with the SEM observations, which showed that the nested porous structures with dimensions of 10^-7^ m or smaller became dominant after 12 h degradation.

Notably, this study revealed several structural changes that occurred specifically in the earlier degradation stage: (i) broadening of the amorphous hallow in WAXD profiles (Figure 2B), (ii) removal of the smooth surface, visualized by SEM (Figures 4 and 5), and (iii) a decrease in the SAXS signal around *q* = 0.022 Å^-1^ (Figure 6A). Summarizing these observations, the surface layer of the amorphous PET film might be slightly more crystalline than the core region and contains the domain (nodule) structure with a size of around 300 Å. This surface layer did not allow PETase to bore smaller holes (∼100 nm) and yielded less porous features, as observed in the SEM images. Thus, this study supports the idea that the surface and core structures differ for amorphous PET films. Feasibly, PETase can be used as a probe to assess the polymer structure of PET in future studies.

### 3.6. Comparison with enzymatic degradation of cellulose

Finally, the results of this study were compared with the structural changes in cellulose degraded by cellulase to discuss a biological strategy for degrading solid polymers. In our previous study [27], the degradation of bacterial cellulose by crude cellulase was analyzed using SAXS, where an overall decrease in the scattering intensity was observed, without changes in the scattering features, indicating that the amount of the cellulose simply decreased without significant structural changes at the scale represented by *q* = 0.04‒0.2 Å^-1^. SEM observations in the same study also demonstrated that the cellulose microfibrils remained generally intact despite the appearance of large cavities between the fiber bundles in a bacterial cellulose film [27]. Another study analyzed the full-width at half-maximum (FWHM) of the diffraction peaks in the WAXD profiles, and also showed that the lateral crystallite size of cellulose decreased because of cellulase degradation [28]. These previous observations of cellulose degradation are in contrast to the structural decay of the PET film by Fast-PETase observed in this study (Figures 2, 5, and 6). This is plausibly due to the structural differences in PET and cellulose due to differences in the assembly of the polymer molecules, as explained below.

Native cellulosic material is generally a woven textile of crystalline cellulose microfibrils with a lateral size of 10^-9^‒10^-8^ m, whereas the PET film is a continuous mass in which many chains are entangled without a well-defined elementary structural unit, such as the microfibrils in cellulose. It is assumed that each cellulose microfibril is degraded independently by cellulase, and the structural features appear to be maintained on average. However, structural degradation of the PET film by Fast-PETase may preferentially occur close to the surface, as revealed by the specific decrease in the SAXS signal at approximately *q* = 0.022 Å^-1^ at the beginning of degradation, which is not observed for cellulose. Therefore, the polymer structure governs the enzymatic degradability. In other words, the design of the polymer structure is as important as enzyme engineering for realizing eco-friendly PET or plastic materials in general.

The relationship between the polymer structure and enzymatic degradation of cellulose has aided in understanding the structure of cellulose and the function of cellulase. For example, the unidirectional degradation of native cellulose microfibrils by cellobiohydrolase was used to demonstrate parallel chain packing in native cellulose microfibrils [29]. In contrast, parallel-chain packing in native cellulose allowed the processive action of cellobiohydrolases to be clearly understood [30–32]. Such interrelated research is also required for PET and PETase to deepen our understanding of the mechanism of decay of the PET structure by PETase. Thus, studying PETase from the viewpoint of the PET structure will be a highly valuable strategy for studying not only PETase but also PET, where many issues remain obscure despite the long history of research [16–18].

Given that cellulase is a good example of the efficient degradation of polymer solids, it will be worthwhile to design a PETase protein by referring to cellulase. The fusion of the carbohydrate-binding module (CBM) to PETase, which is also found in nature [33], has been shown to enhance the PET-degrading activity [34–37]. In addition to implementing the absorption function by CBM, the synergistic effect of mixing enzymes with different modes of action, such as exotype cellulases (cellobiohydrolases) and endoglucanases, is a promising strategy [23]. To this end, the mode of action of many PETase proteins must be clarified to design a high-performance PETase cocktail. A cellulase-wise strategy is a promising way to develop higher-performance PETases. This approach is set apart from state-of-the-art technologies, such as protein engineering with bioinformatics. Polymer structure analysis during enzymatic degradation is important for the sustainable utilization of plastic materials and for the research and development of biodegradable plastics.

## 4. Conclusion

The multiscale structural analysis in this study revealed that Fast-PETase gradually eroded the PET film from the surface and bored many pores with diameters of 10^-5^ to 10^-8^ m. The domain structure with dimensions of 10^-8^ m on the surface of the film was preferentially removed at earlier reaction times. However, the crystallinity of PET did not change after degradation. In summary, it was demonstrated that Fast-PETase evenly erodes the amorphous PET film from the surface. This study provides a perspective on the degradation of the solid PET by PETase. This is valuable information as a reference for comparing the enzymatic efficacy of PETase variants for PET degradation that are still to be discovered and will help pave the way for a circular economy of PET materials in the future by conferring full biodegradability to PET, a conventional plastic.

## Acknowledgments

SEM observations were performed at a collaborative facility at the Research Institute for Sustainable Humanosphere (RISH) at Kyoto University, Cellulosic Advanced Nanomaterials Development Organization (CAN-DO). The micro-X-ray CT experiment was performed at BL8S2 at Aichi SRC (202403106). SAXS measurements were performed at BL40B2 (2023B1470, 2024A1381, and 2024B1389) and BL19B2 (2024A1728) at SPring-8 and BL8S3 at Aichi SRC (202306168). The WAXD measurements were performed at BL40B2 at SPring-8 (2023B1470, 2024A1381, and 2024B1389)). We thank the beamline staff at the synchrotron facilities for their supporting the experiments. We acknowledge Dr. Susumu Fujiwara in Kyoto Institute of Technology, Dr. Shosuke Yoshida in Nara Institute of Science and Technology, and Dr. Takashi Konishi in Kyoto University for discussing the experimental results.

## Author contributions

DT conceptualization; DT and TI methodology; DT and TI formal analysis; DT and TI investigation; DT and TI resources; DT and TI data curation; DT and TI writing–original draft; TI writing–review and editing; DT and TI visualization; TI supervision; TI project administration; DT and TI funding acquisition.

## References

[1] V. Hidalgo-Ruz, L. Gutow, R.C. Thompson, M. Thiel, Microplastics in the Marine Environment: A Review of the Methods Used for Identification and Quantification, Environ. Sci. Technol. 46(6) (2012) 3060–3075.

[2] A.A. Horton, Plastic Pollution in the Global Ocean, WORLD SCIENTIFIC, Singapore, 2022.

[3] Y. Li, L. Chen, N. Zhou, Y. Chen, Z. Ling, P. Xiang, Microplastics in the human body: A comprehensive review of exposure, distribution, migration mechanisms, and toxicity, Sci. Total Environ. 946 (2024) 174215.

[4] K. Ragaert, L. Delva, K. Van Geem, Mechanical and chemical recycling of solid plastic waste, Waste Manag. 69 (2017) 24–58.

[5] C. Jehanno, J.W. Alty, M. Roosen, S. De Meester, A.P. Dove, E.Y.X. Chen, F.A. Leibfarth, H. Sardon, Critical advances and future opportunities in upcycling commodity polymers, Nature 603(7903) (2022) 803–814.

[6] M. Tsuchiya, T. Kitahashi, R. Nakajima, K. Oguri, K. Kawamura, A. Nakamura, K. Nakano, Y. Maeda, M. Murayama, S. Chiba, K. Fujikura, Distribution of microplastics in bathyal- to hadal-depth sediments and transport process along the deep-sea canyon and the Kuroshio Extension in the Northwest Pacific, Mar. Pollut. Bull. 199 (2024) 115466.

[7] M.H. Ghasemi, N. Neekzad, F.B. Ajdari, E. Kowsari, S. Ramakrishna, Mechanistic aspects of poly(ethylene terephthalate) recycling–toward enabling high quality sustainability decisions in waste management, Environ. Sci. Pollut. Res. 28(32) (2021) 43074–43101.

[8] R.-J. Müller, H. Schrader, J. Profe, K. Dresler, W.-D. Deckwer, Enzymatic Degradation of Poly(ethylene terephthalate): Rapid Hydrolyse using a Hydrolase from T. fusca, Macromol. Rapid Commun. 26(17) (2005) 1400–1405.

[9] S. Yoshida, K. Hiraga, T. Takehana, I. Taniguchi, H. Yamaji, Y. Maeda, K. Toyohara, K. Miyamoto, Y. Kimura, K. Oda, A bacterium that degrades and assimilates poly(ethylene terephthalate), Science 351(6278) (2016) 1196–1199.

[10] S. Sulaiman, S. Yamato, E. Kanaya, J.-J. Kim, Y. Koga, K. Takano, S. Kanaya, Isolation of a Novel Cutinase Homolog with Polyethylene Terephthalate-Degrading Activity from Leaf-Branch Compost by Using a Metagenomic Approach, Appl. Environ. Microbiol. 78(5) (2012) 1556–1562.

[11] H. Lu, D.J. Diaz, N.J. Czarnecki, C. Zhu, W. Kim, R. Shroff, D.J. Acosta, B.R. Alexander, H.O. Cole, Y. Zhang, N.A. Lynd, A.D. Ellington, H.S. Alper, Machine learning-aided engineering of hydrolases for PET depolymerization, Nature 604(7907) (2022) 662–667.

[12] S.H. Lee, H. Seo, H. Hong, J. Park, D. Ki, M. Kim, H.-J. Kim, K.-J. Kim, Three-directional engineering of IsPETase with enhanced protein yield, activity, and durability, J. Hazard. Mater. 459 (2023) 132297.

[13] S. Joo, I.J. Cho, H. Seo, H.F. Son, H.-Y. Sagong, T.J. Shin, S.Y. Choi, S.Y. Lee, K.-J. Kim, Structural insight into molecular mechanism of poly(ethylene terephthalate) degradation, Nature Communications 9(1) (2018) 382.

[14] S. Weigert, P. Perez-Garcia, F.J. Gisdon, A. Gagsteiger, K. Schweinshaut, G.M. Ullmann, J. Chow, W.R. Streit, B. Höcker, Investigation of the halophilic PET hydrolase PET6 from Vibrio gazogenes, Protein Sci. 31(12) (2022) e4500.

[15] T. Burgin, B.C. Pollard, B.C. Knott, H.B. Mayes, M.F. Crowley, J.E. McGeehan, G.T. Beckham, H.L. Woodcock, The reaction mechanism of the Ideonella sakaiensis PETase enzyme, Commun. Chem. 7(1) (2024) 65.

[16] P.H. Geil, Morphology of Amorphous Polymers, Product R&D 14(1) (1975) 59–71.

[17] A.L. Renninger, D.R. Uhlmann, On the structure of glassy polymers. III. Small-angle x-ray scattering from amorphous polyethylene terephthalate, J. Polym. Sci.: Polym. Phys. Ed. 14(3) (1976) 415–425.

[18] P. Calvert, Order in amorphous polymers, Nature 271(5645) (1978) 507–508.

[19] F. Teufel, J.J.A. Armenteros, A.R. Johansen, M.H. Gíslason, S.I. Pihl, K.D. Tsirigos, O. Winther, S. Brunak, G. von Heijne, H. Nielsen, SignalP 6.0 predicts all five types of signal peptides using protein language models, Nat. Biotechnol. 40(7) (2022) 1023–+.

[20] D.a. Gürsoy, F. De Carlo, X. Xiao, C. Jacobsen, TomoPy: a framework for the analysis of synchrotron tomographic data, J. Synchrotron Radiat. 21(5) (2014) 1188–1193.

[21] J. Schindelin, I. Arganda-Carreras, E. Frise, V. Kaynig, M. Longair, T. Pietzsch, S. Preibisch, C. Rueden, S. Saalfeld, B. Schmid, J.-Y. Tinevez, D.J. White, V. Hartenstein, K. Eliceiri, P. Tomancak, A. Cardona, Fiji: an open-source platform for biological-image analysis, Nat. Methods 9(7) (2012) 676–682.

[22] K. Tanaka, A. Takahara, T. Kajiyama, Rheological Analysis of Surface Relaxation Process of Monodisperse Polystyrene Films, Macromolecules 33(20) (2000) 7588–7593.

[23] T.T. Teeri, Crystalline cellulose degradation: new insight into the function of cellobiohydrolases, Trends Biotechnol. 15(5) (1997) 160–167.

[24] C.M. Payne, B.C. Knott, H.B. Mayes, H. Hansson, M.E. Himmel, M. Sandgren, J. Ståhlberg, G.T. Beckham, Fungal Cellulases, Chem. Rev. 115(3) (2015) 1308–1448.

[25] G.S.Y. Yeh, Morphology of amorphous polymers and effects of thermal and mechanical treatments on the morphology, Pure Appl. Chem. 31(1-2) (1972) 65–90.

[26] T. Konishi, D. Okamoto, D. Tadokoro, Y. Kawahara, K. Fukao, Y. Miyamoto, Origin of SAXS intensity in the low-q region during the early stage of polymer crystallization from both the melt and glassy state, Phys. Rev. Mater. 2(10) (2018) 105602.

[27] P.A. Penttilä, T. Imai, J. Hemming, S. Willför, J. Sugiyama, Enzymatic hydrolysis of biomimetic bacterial cellulose¥textendashhemicellulose composites, Carbohydr. Polym. 190 (2018) 95–102.

[28] T. Imai, M. Naruse, Y. Horikawa, K. Yaoi, K. Miyazaki, J. Sugiyama, Disturbance of the hydrogen bonding in cellulose by bacterial expansin, Cellulose 30 (2023) 8423–8438.

[29] H. Chanzy, B. Henrissat, Undirectional degradation of valonia cellulose microcrystals subjected to cellulase action, FEBS Lett. 184(2) (1985) 285–288.

[30] T. Imai, C. Boisset, M. Samejima, K. Igarashi, J. Sugiyama, Unidirectional processive action of cellobiohydrolase Cel7A on Valonia cellulose microcrystals, FEBS Lett. 432(3) (1998) 113–116.

[31] N.-H. Kim, T. Imai, M. Wada, J. Sugiyama, Molecular directionality in cellulose polymorphs, Biomacromolecules 7(1) (2006) 274–280.

[32] K. Igarashi, A. Koivula, M. Wada, S. Kimura, M. Penttila, M. Samejima, High speed atomic force microscopy visualizes processive movement of Trichoderma reesei cellobiohydrolase I on crystalline cellulose, J. Biol. Chem. 284(52) (2009) 36186–36190.

[33] E. Erickson, J.E. Gado, L. Avilán, F. Bratti, R.K. Brizendine, P.A. Cox, R. Gill, R. Graham, D.-J. Kim, G. König, W.E. Michener, S. Poudel, K.J. Ramirez, T.J. Shakespeare, M. Zahn, E.S. Boyd, C.M. Payne, J.L. DuBois, A.R. Pickford, G.T. Beckham, J.E. McGeehan, Sourcing thermotolerant poly(ethylene terephthalate) hydrolase scaffolds from natural diversity, Nat. Commun. 13(1) (2022) 7850.

[34] Y. Zhang, L. Wang, J. Chen, J. Wu, Enhanced activity toward PET by site-directed mutagenesis of Thermobifida fusca cutinase–CBM fusion protein, Carbohydr. Polym. 97(1) (2013) 124–129.

[35] J. Weber, D. Petrovic, B. Strodel, S.H.J. Smits, S. Kolkenbrock, C. Leggewie, K.E. Jaeger, Interaction of carbohydrate-binding modules with poly(ethylene terephthalate), Appl. Microbiol. Biotechnol. 103(12) (2019) 4801–4812.

[36] L. Dai, Y. Qu, J.-W. Huang, Y. Hu, H. Hu, S. Li, C.-C. Chen, R.-T. Guo, Enhancing PET hydrolytic enzyme activity by fusion of the cellulose–binding domain of cellobiohydrolase I from Trichoderma reesei, J. Biotechnol. 334 (2021) 47–50.

[37] R. Graham, E. Erickson, R.K. Brizendine, D. Salvachúa, W.E. Michener, Y. Li, Z. Tan, G.T. Beckham, J.E. McGeehan, A.R. Pickford, The role of binding modules in enzymatic poly(ethylene terephthalate) hydrolysis at high-solids loadings, Chem Catalysis 2(10) (2022) 2644–2657.

